# Connectome spectral analysis to track EEG task dynamics on a subsecond scale

**DOI:** 10.1101/2020.06.22.164111

**Authors:** Katharina Glomb, Joan Rue Queralt, David Pascucci, Michaël Defferrard, Sebastien Tourbier, Margherita Carboni, Maria Rubega, Serge Vulliemoz, Gijs Plomp, Patric Hagmann

## Abstract

We present an approach for tracking fast spatiotemporal cortical dynamics in which we combine white matter connectivity data with source-projected electroencephalographic (EEG) data. We employ the mathematical framework of *graph signal processing* in order to derive the Fourier modes of the brain structural connectivity graph, or “network harmonics”. These network harmonics are naturally ordered by smoothness. Smoothness in this context can be understood as the amount of variation along the cortex, leading to a multi-scale representation of brain connectivity. We demonstrate that network harmonics provide a sparse representation of the EEG signal, where, at certain times, the smoothest 15 network harmonics capture 90% of the signal power. This suggests that network harmonics are functionally meaningful, which we demonstrate by using them as a basis for the functional EEG data recorded from a face detection task. There, only 13 network harmonics are sufficient to track the large-scale cortical activity during the processing of the stimuli with a 50 ms resolution, reproducing well-known activity in the fusiform face area as well as revealing co-activation patterns in somatosensory/motor and frontal cortices that an unconstrained ROI-by-ROI analysis fails to capture. The proposed approach is simple and fast, provides a means of integration of multimodal datasets, and is tied to a theoretical framework in mathematics and physics. Thus, network harmonics point towards promising research directions both theoretically - for example in exploring the relationship between structure and function in the brain - and practically - for example for network tracking in different tasks and groups of individuals, such as patients.

## 1. Introduction

Many recent studies in neuroscience have stressed the importance of spatiotemporal dynamics of brain activity for our understanding of brain function (Cichy et al. 2016; Northoff, Wainio-Theberge, and Evers 2019; Deco et al. 2019; Atasoy, Deco, and Kringelbach 2019). Electroencephalography (EEG) records brain activity with millisecond temporal resolution, allowing, in principle, tracking of brain dynamics on a behaviorally relevant time scale. Furthermore, EEG is portable and relatively cheap compared to functional magnetic resonance imaging (fMRI) or magnetoencephalography (MEG), and therefore has large clinical potential. Approaches that use functional magnetic resonance imaging (fMRI) data have revealed many important insights into (ultra-) slow network dynamics (Deco et al. 2013; Tagliazucchi et al. 2012; Griffa et al. 2017; Karahanoğlu and Van De Ville 2015; Preti, Bolton, and Van De Ville 2017; Lurie et al. 2020), and several approaches tackle tracking of fast network dynamics using specifically M/EEG activity projected into the gray matter source space (Silfverhuth et al. 2012; Mheich et al. 2015; Quinn et al. 2018; Kabbara et al. 2019; Baker et al. 2014; Tewarie, Liuzzi, et al. 2019; Zerouali et al. 2014; Coito et al. 2016; Brookes et al. 2014). These approaches typically derive functional networks directly from the functional data. The issue is that EEG activity is highly impacted by the effects of volume conduction and the regularization performed in the lead field models, producing a correlation structure among the sources that is determined by Euclidean distance between sources beyond the real functional connectivity that exists between nearby brain regions. This is true even when measures of functional connectivity are used that reduce this impact (O’Neill et al. 2018), for example, imaginary coherence (Nolte et al. 2004) or phase lag index (Stam et al. 2007), both of which remove zero lag phase interactions between signals, or if functional connectivity is computed after linear dependencies between signals are removed via orthogonalization (Brookes, Woolrich, and Barnes 2012; Colclough et al. 2015)

In this study we propose to use Fourier modes extracted from the brain structural connectivity graph, or connectome, i.e. network harmonics, as an alternative approach. It takes advantage of different data modalities - functional and structural - and combines them in a mathematically principled way that is rooted in theory and connects this approach to physics applications by performing a graph spectral analysis. In this view, brain regions, as defined by an atlas/a parcellation, form nodes of a graph which are linked by edges whose weights depend on, in this case, white matter anatomical connections. Network harmonics optimally preserve the local graph structure on different scales (Belkin and Niyogi 2003), which means that they minimize differences between neighboring nodes in the graph. This way, they provide building blocks of functional activity in which anatomically connected regions co-activate, which has been shown to result in more efficient switching between patterns (Gu et al. 2018; Tomasi, Wang, and Volkow 2013). Importantly, in network harmonics, two brain regions, or nodes, are not simply assigned to *either* the same network *or* to different networks, but the degree to which their connectivity patterns to the rest of the graph resemble each other is taken into account. Thus, a network harmonic can be interpreted as a connectivity gradient (Margulies et al. 2016; Haak, Marquand, and Beckmann 2018; Atasoy, Donnelly, and Pearson 2016; Glomb, Kringelbach, et al. 2019).

Mathematically, network harmonics are computed as the eigenvectors of the graph Laplacian of the structural connectivity (Shuman et al. 2012; Belkin and Niyogi 2003). Put another way, network harmonics are the basis functions of the brain network encoded by the structural connectivity matrix. This is equivalent to sine and cosine waves being basis functions of the unit circle: superpositions of these basis functions approximate any signal on their domain (the graph or the circle, respectively) to arbitrary precision (Atasoy et al. 2017), which is the idea underlying Fourier series. In continuous domains, basis functions of the Laplace operators are used in many physical applications, for example pattern formation (Murray 1988), or the famous example of Chladni’s vibrating metal plates (Leissa 1973). (Robinson et al. 2016) have shown that applying spherical harmonics to the brain can explain important features of large-scale networks from the point of view of neural field theory.

In this study, we consider EEG activity in the framework of graph signal processing, i.e. as a signal on the domain of the “brain graph”, and as such, it can be described by a superposition of the connectivity gradients encoded in network harmonics. This amounts to a graph (spatial) spectral representation of the EEG signal. Quantifying this spectrum over time allows to track fast dynamics of large-scale networks.

The idea of network harmonics allows the direct integration of different sources of data in a single graph. Specifically, the graph mentioned above, and the network harmonics themselves, are derived from diffusion MRI, while the functional data were recorded with EEG. It has been shown that in the domain of network harmonics, structural and functional connectivity are analytically related, both in fMRI (Abdelnour et al. 2018; Atasoy, Donnelly, and Pearson 2016; Raj et al. 2020) and MEG (Tewarie, Abeysuriya, et al. 2019). More generally, SC shapes functional activity, as has been shown for EEG (Chu et al. 2015; Wirsich et al. 2017; Finger et al. 2016; Glomb, Mullier, et al. 2019), MEG (Cabral et al. 2014), and fMRI (Deco et al. 2013). In this paper, we apply the mathematical tools of graph signal processing to a multimodal dataset consisting of structural, functional, and diffusion data, extracting network harmonics as building blocks of large-scale cortical activity. We first demonstrate that network harmonics provide a sparser representation of source-level task EEG data than the region-by-region representation. We find that most of the signal is captured by the smoothest network harmonics. We show that this framework, when used to analyze data recorded during a facial recognition task, reveals fast dynamics of large-scale networks associated with the processing of such stimuli. This framework offers a mathematically principled and multimodal approach to fast spatiotemporal network dynamics in EEG.

## 2. Methods

### 2.1. EEG Data acquisition and preprocessing

128 channel high-density EEG was recorded at 2048 Hz with a 128-channel Biosemi Active Two EEG system (Biosemi, Amsterdam, The Netherlands; details on the EEG montage [electrode positions] can be downloaded from the manufacturer’s website at www.biosemi.com/download/ → Cap_coords_all.xls) at Hopital Cantonal Fribourg from 20 healthy participants (17 females, mean age: 23, age range: 20-29 years) performing a visual discrimination task. During recording, good signal quality was guaranteed by keeping the offset between the active electrodes and the Common Mode Sense - Driven Right Leg (CMS-DRL) feedback loop under a standard value of ±20 mV. After each recording session, individual 3D electrode positions were digitized using an ultrasound motion capture system (Zebris Medical GmbH). A subset of these data have been used in a previous publication (Rubega et al. 2019). Participants viewed images of faces or scrambled versions of the same images (Ales et al. 2012), presented for 200 ms, and then responded, by pressing one of two buttons on a response box, whether they had seen a face or a scrambled image. One participant was excluded due to too many motion artifacts, leaving 19 datasets for analysis. Data were preprocessed using EEGLAB (Delorme and Makeig 2004). EEGLAB is freely available for download at sccn.ucsd.edu/eeglab/index.php. First, the time series were downsampled to 250 Hz (anti-aliasing filter: cut-off frequency = 112.5 Hz; transition bandwidth = 50 Hz) and local detrending (high-pass filter at 1 Hz, EEGLAB PREP plugin) was applied (Bigdely-Shamlo et al. 2015). Epochs from 1500 ms before until 1000 ms after stimulus onset were extracted. Line and monitor noise (at 50 and 75 Hz, respectively, as well as harmonics of these frequencies) were removed by spectral interpolation (Leske and Dalal 2019). Bad epochs (22 ± 36 out of 600 per subject) were removed and bad channels (15 ± 9 out of 128 per subject) marked via visual inspection. Next, remaining physiological artifacts (eye blinks, horizontal and vertical eye movements, muscle potentials) were removed using FastICA by first marking bad ICs using Multiple Artifact Rejection Algorithm (MARA) as implemented in EEGLAB (Delorme and Makeig 2004). The previously identified bad channels were not included in this step. Finally, bad channels were interpolated using the nearest neighbor spline method as implemented in EEGLAB, and data were re-referenced to the common average before being globally z-scored.

### 2.2. Structural images and tissue segmentation

T1-weighted images obtained from the same subjects was acquired as magnetization prepared rapid-gradient echo (MPRAGE) volumes with a General Electrics Discovery MR750 3T MRI scanner and a COR FSPGR BRAVO pulse sequence with flip angle = 9°; echo time = 2.81 ms, repetition time = 7.27 ms, inversion time = 0.9 s, slice thickness = 1 mm, head first supine. Segmentation of the MPRAGE volume into gray and white matter was performed using Connectome mapper 3 (Tourbier et al. 2020) with Freesurfer 6.0.1, and gray matter brain regions were subsequently extracted according to the Lausanne 2008 multiscale parcellation (Hagmann et al. 2008). Connectome Mapper 3 and the Lausanne multiscale parcellation are available at github.com/connectomicslab/connectomemapper3. Freesurfer can be downloaded from surfer.nmr.mgh.harvard.edu/fswiki/FreeSurferWiki.

### 2.3. Source projection and ROI-time course extraction

In order to project EEG signals recorded on the scalp into the gray matter, individual head models were created (Brunet, Murray, and Michel 2011) based on the structural images and segmented tissues explained above.

Inverse solutions were computed using LAURA (Local Autoregressive Average) with LSMAC (Locally Spherical Model with Anatomical Constraints) as implemented in CARTOOL (Brunet et al. 2011; Grave de Peralta Menendez et al. 2004). LSMAC is based on a 3-shell head model that takes into account the local radiuses of the skull, scalp, and brain, and is simpler and therefore computationally less expensive than boundary of finite element models (BEM/FEM, respectively; (Birot et al. 2014)). It uses a regularization technique based on models of the electrical generators of the EEG signal (Grave de Peralta Menendez et al. 2004), and the best regularization parameter is chosen according to the L-curve (Hansen 1992). CARTOOL is freely available for download at sites.google.com/site/cartoolcommunity.

Data recorded on the scalp, i.e. in sensor space, were projected to ~5000 dipole locations equally spaced on a 3D-dimensional-grid within each individual’s gray matter, i.e. into source space. (This gray matter volume, as well as the other tissue types needed for the individual head model, were extracted from the structural scans described above.)

The dipole time courses are three-dimensional as they have a magnitude and a direction in xyz-space, and thus, one-dimensional time courses for further analysis have to be extracted. We use singular value decomposition as described in (Rubega et al. 2019)First, the main direction of variance for each individual dipole time course is extracted by averaging over all epochs within a subject and performing singular value decomposition on the matrix of average activity time courses for a time window of interest, here, 100 to 300 ms after stimulus onset, where the strongest response is expected. The first left-singular vector is the direction of greatest variance. The single trial time courses of each dipole are then projected onto this direction, preserving most of the variance. Signs of the first left-singular vectors were aligned across trials and subjects. Finally, the thus obtained 1-dimensional source time courses are averaged within each brain region defined by the individual parcellation in subject space. Importantly, even though parcellations and inverse solutions are based on individual MRI data, and are performed in individual space (i.e. not transformed into a standardized template space), one-dimensional region time courses can be averaged across participants, assuming correspondence between the atlas’s regions. Note that in principle, the unit of the signal is the same in source space as in sensor space, i.e. micro-volts. However, due to the many steps in forward modelling, projecting, and averaging, we merely refer to the signal as “activity” or “amplitude” and use arbitrary units (a.u.) throughout the paper.

### 2.4. Consensus structural connectivity matrix

We use a consensus connectome from 88 healthy participants (70 of which are available online, (Griffa, Alemán-Gómez, and Hagmann 2019); mean age 29.7 years, minimum 18.5, maximum 59.2 years; 34 females), scanned in a 3-Tesla MRI scanner (Trio, Siemens Medical, Germany) with a 32-channel head-coil. All subjects are part of an ongoing study on schizophrenia at Centre Hospitalier Universitaire Vaudois (CHUV) in Lausanne, Switzerland, and the 18 subjects whose data are not available online were recorded and preprocessed after the release of the dataset. A diffusion spectrum imaging (DSI) sequence (128 diffusion-weighted volumes and a single b0 volume, maximum b-value 8,000 s/mm^2^, 2.2×2.2×3.0 mm voxel size) was applied, and DSI data were reconstructed following the protocol described in (Wedeen et al. 2005). In brief, multiple diffusion directions per voxel were estimated, reconstructing the diffusion probability density function as the discrete 3D Fourier transform of the signal modulus. The 3D probability distribution function was then normalized, and the orientation distribution function (ODF) computed as the radial summation of the normalized probability distribution function. Thus, the ODF is defined on a discrete sphere and captures the diffusion intensity in every direction. A magnetization-prepared rapid acquisition gradient echo (MPRAGE) sequence sensitive to white/gray matter contrast (1-mm in-plane resolution, 1.2-mm slice thickness) was also acquired, and gray and white matter were segmented from the MPRAGE volume using Freesurfer and Connectome Mapper 3 (Tourbier et al. 2020).

Individual structural connectivity matrices were estimated using deterministic streamline tractography on reconstructed DSI data, initiating 32 streamline propagations per diffusion direction, per white matter voxel (Wedeen et al. 2008). The number of fibers found between each voxel at the gray matter/white matter-interface was summed within each ROI given by the same parcellation used above for the structural and functional data (Desikan et al. 2006). Matrices were subsequently averaged across subjects. Note that no morphing into standard space was necessary, as we averaged using the atlas parcels. We applied the method introduced in (Betzel et al. 2019). Briefly, this method selects a recurrence threshold (i.e. a threshold on how many subjects are required to have a non-zero fiber count between a given pair of brain regions) which preserves the connection density of single subject SCs for intra- and interhemispheric connections separately. By doing so, a less conservative threshold is applied to the interhemispheric connections, as they are harder to track and are thus less reproducible across subjects. The resulting connection density in our SC is 25%.

### 2.5. Network harmonics and graph visualization

The basic idea is that co-activation patterns on the connectivity graph that describes the brain can be used as “building blocks” of brain activity on the same domain. These building blocks are obtained as the harmonic basis functions, or harmonic modes, of the structural connectivity, i.e. by performing an eigenvalue decomposition of the SC’s graph Laplacian. In these basis functions, local distances on the graph are preserved (Belkin and Niyogi 2003), such that strongly connected pairs of brain regions exhibit similar values in the eigenvectors (see also Figure 1c). When interpreting the eigenvectors as building blocks of brain activity, this preservation of structural connections results in connected nodes being co-activated. The graph Laplacian is the graph equivalent of the second spatial derivative (i.e., the Laplace operator) in the continuous domain, whose eigenvectors are well-known in some cases: in the case of the circular domain, they form the Fourier basis functions (i.e., sine and cosine waves), spherical harmonics in the case of the sphere (Robinson et al. 2016).

**Figure 1.**
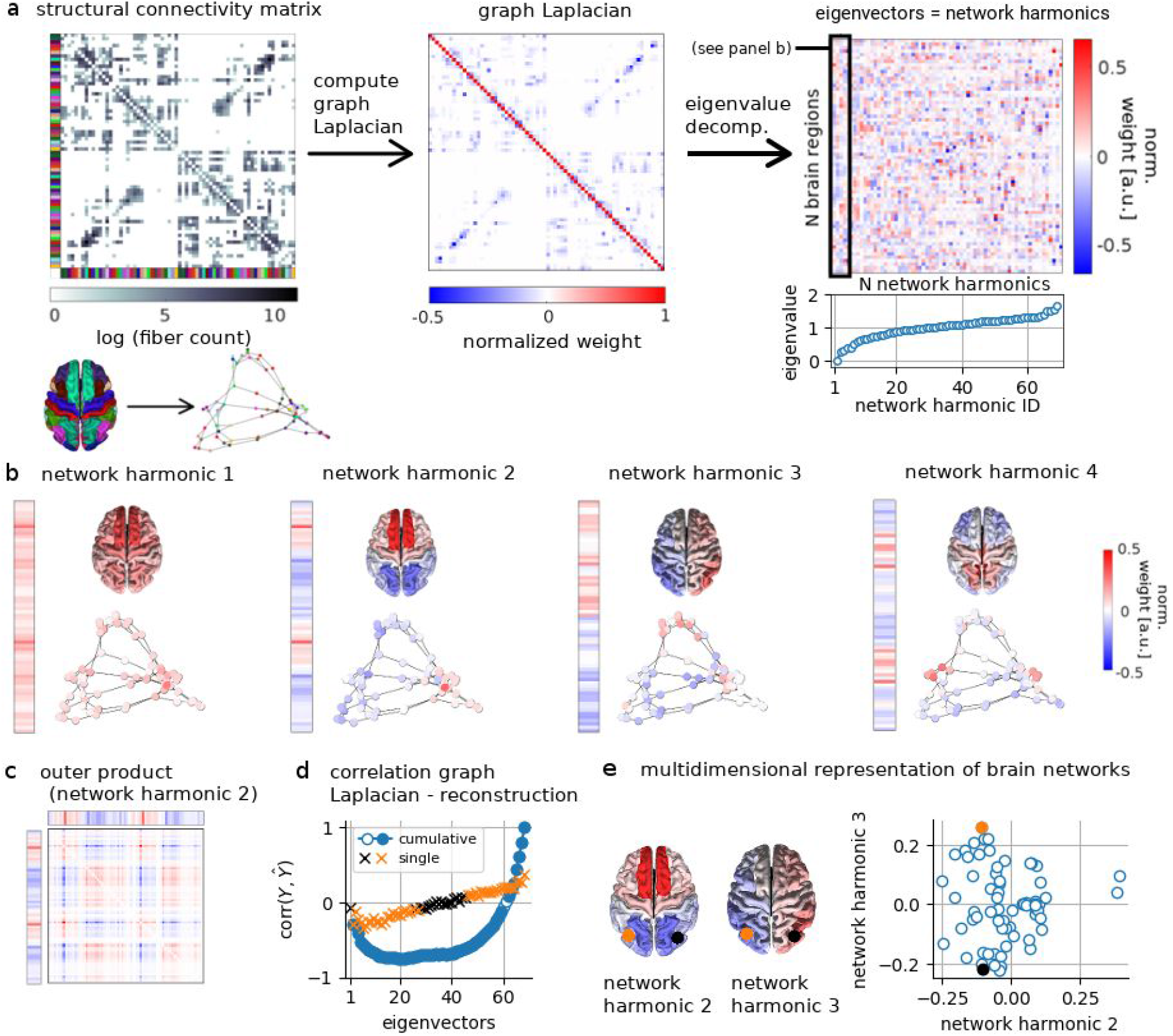
**a:** Illustration of workflow, from left to right. The structural connectivity matrix encodes a graph in which each node is a brain region defined by a parcellation (see brain surface below matrix; brain regions, nodes, and matrix rows/columns are color-coded) and edges are defined by white matter connectivity. The graph Laplacian is computed from this matrix, and network harmonics are obtained as eigenvectors of this graph Laplacian. The eigenvectors are ordered by ascending eigenvalue. **b:** The first four network harmonics in vector form (magnified from panel **a**), projected onto the surface of the brain, and in graph representation. Colors visualize arbitrary units, i.e. the weights in the orthonormal eigenvectors. **c:** The graph Laplacian can be reconstructed as a weighted sum of rank-1 matrices defined by the outer products of its eigenvectors. Large values of equal sign in the eigenvector (network harmonic) lead to large positive weights in the outer product (red entries in the illustration). **d**: The correlation between the (upper or lower triangle) of the reconstructed matrices 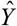 obtained from the outer products and the original graph Laplacian *Y* is used to quantify how well the eigenvectors capture the SC, both when they are used cumulatively (circles) and on their own (crosses). Open blue circles and black crosses marknon-significant correlations for each case. **e**: Two brain regions, here: inferior parietal, are close together in network harmonic 2 (x-axis), but far apart in network harmonic 3 (y-axis), illustrating how network harmonics capture integration and segregation in multiple dimensions.

The network harmonics used in this paper are computed as the eigenvectors of the normalized graph Laplacian. We compute the normalized graph Laplacian *L* from the SC *C* as follows:

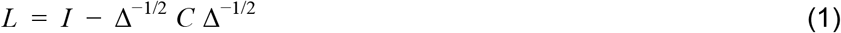

Δ is the degree matrix, *I* is the identity matrix. The resulting matrix *L*, the normalized graph Laplacian, has ones in its diagonal and negative entries for pairs that have non-zeros connectivity in *C*. Each row and column sums to 0. *L* is then decomposed into eigenvectors and eigenvalues. The resulting eigenvectors, or eigenfunctions, form a set of orthonormal basis functions just like sine and cosine waves do in the “conventional” Fourier transform. The first 4 eigenvectors are shown in Figure 1b and the first 10 in Figure S1. The i-th eigenvalue is a measure of the smoothness of the i-th eigenvector because

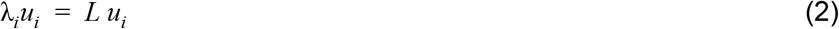

per definition of the eigendecomposition *L* = *U* Λ*U^T^* with eigenvectors *u*_*i*_ such that *U* = [*u*_1_, *u*_2_, …, *u_*N*_*] and eigenvalues λ_*i*_ are entries of the diagonal matrix Λ = *diag*(λ_1_, …, λ◻). Then

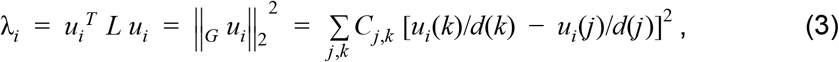

where _*G*_ is the gradient operator on the graph and 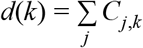 is the weighted degree.

Hence, λ_*i*_ is a measure of how much *u*_*i*_ varies across connected vertices. This quadratic form is known as the *Dirichlet energy* of the signal *u*_*i*_. For more details, please refer to (Shuman et al. 2012).

The graph Laplacian can be approximated using only a subset of the eigenvectors and -values:

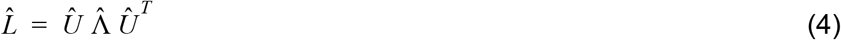

Here, 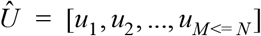 and, 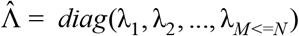 or correspondingly for single eigenvectors and -values, 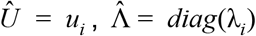 In order to quantify how well each sequence of eigenvectors or each single eigenvector captures the brain network, we compute the correlation between the entries which have non-zero values in the Laplacian (existing connections) - denoted by *Y* - and the same entries in the reconstruction, denoted by. 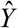

The graph representation shown in Figure 1 was obtained using networkx’ spectral_layout() function, which returns node positions based on the first two Laplace eigenvectors. The toolbox including documentation is available on networkx.github.io/documentation/stable/index.html. Randomized graphs were obtained using the Brain Connectivity Toolbox (Rubinov and Sporns 2010) function “null_model_und_sign.m”. The Brain Connectivity Toolbox can be freely downloaded from sites.google.com/site/bctnet/.

### 2.6. Graph Fourier transform

The EEG time series of each subject is an array of dimensionality *N × T × E*, where *N* is the number of brain regions (*N* = 68), *T* is the number of time points (*T* = 625 for 2.5 s of data with sampling frequency of 250 Hz), and *E* is the number of epochs (or trials), which varies across participants. Each *N* × 1 column of this array is an activation pattern 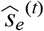 indexed by epoch/trial *e* and time (*t*) ; activation pattern here refers to the amplitude of the one-dimensional ROI signal extracted as explained above, i.e. the electrical activity projected into the gray matter. We refer to this representation of the signal as the “original” domain. The activation patterns are transformed into the space of the network harmonics (i.e., the eigenvectors derived from the normalized graph Laplacian as described above), a transformation that is known as graph Fourier transform (GFT) in the graph signal processing (GSP) literature (Shuman et al. 2012):

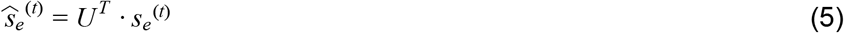

Here, *U* is the *N* x *N* matrix which contains in its columns the network harmonics/eigenvectors of the graph Laplacian, the superscript *T* marks the transpose (*U* is always real-valued), and · denotes the dot product. *s_e_*^(*t*)^ is an activation pattern. 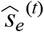 is therefore an *N* x 1 vector containing GFT weights that quantify the contribution of each network harmonic to this particular pattern, or in other words, its graph Fourier spectrum. We refer to this representation of the signal as “graph frequency” or “spectral” domain. Note that the “original” domain in terms of N brain regions is a representation of the signal in space and time, and by applying the GFT, we perform a Fourier transformation along the *spatial* dimension. In principle, a classical Fourier transform could simultaneously be applied along the temporal dimension. In both cases, this transformation constitutes a change of basis, as no information is lost. For the GFT along the spatial domain as used here, the original signal can be reconstructed using the inverse GFT:

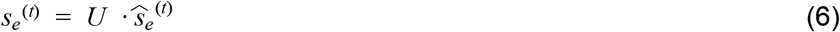

Here, *U* can be substituted by 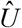 in a similar way as in equation (4). Arbitrary sets of network harmonics (eigenvectors) can be used, resulting in a “reduced”, or “filtered”, reconstructed signal. In this study, we apply this approach when reconstructing the EEG signal in the “original” domain using only a subset of network harmonics that we identify has significantly contributing to the signal, as described below in section “*Statistical analysis of network harmonic activations (sliding window analysis)*”.

### 2.7. Statistical analysis of L1 norm

For the analysis of the sparsity of the signal, we tested how well the representation of the signal in each of the two domains - i.e. “original” domain, N brain regions, and graph frequency, or spectral, domain, N network harmonics (see Figure 2a) captures the difference between pre- and post-stimulus intervals. We picked 50 time points (200 ms) immediately preceding stimulus onset (−200 to 0 ms), and an interval around the peak inflection following the stimulus (140 to 340 ms). We computed the difference vectors between the two intervals (post minus pre) in both domains (“original” domain:difference between average amplitude of N=68 brain regions during these intervals; spectral domain: GFT weights of the network harmonics: difference between average GFT weights of N=68 network harmonics during these intervals). We computed the average distance vector after normalizing each individual’s vector’s power (L2 norm) to 1 in order to remove differences in overall power and focus on the pattern of activation differences. We computed the L1 norm of the average distance vectors for each domain, i.e. the sum of the absolute values of all entries of the vector. In order to evaluate whether the difference between the L1 norms was significant, we conducted a permutation test with n=10.000 permutations, in which we switched the difference vectors between the two domains for a randomly selected subset of the subjects. Thus, we re-computed the average difference vectors from sets of individual difference vectors which contained vectors from both domains. We computed the p-value as

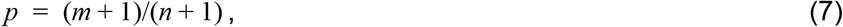

where n is the number of permutations and m is the number of permutations in which the difference between the L1 norms computed from permuted distances exceeded the empirical one.

**Figure 2.**
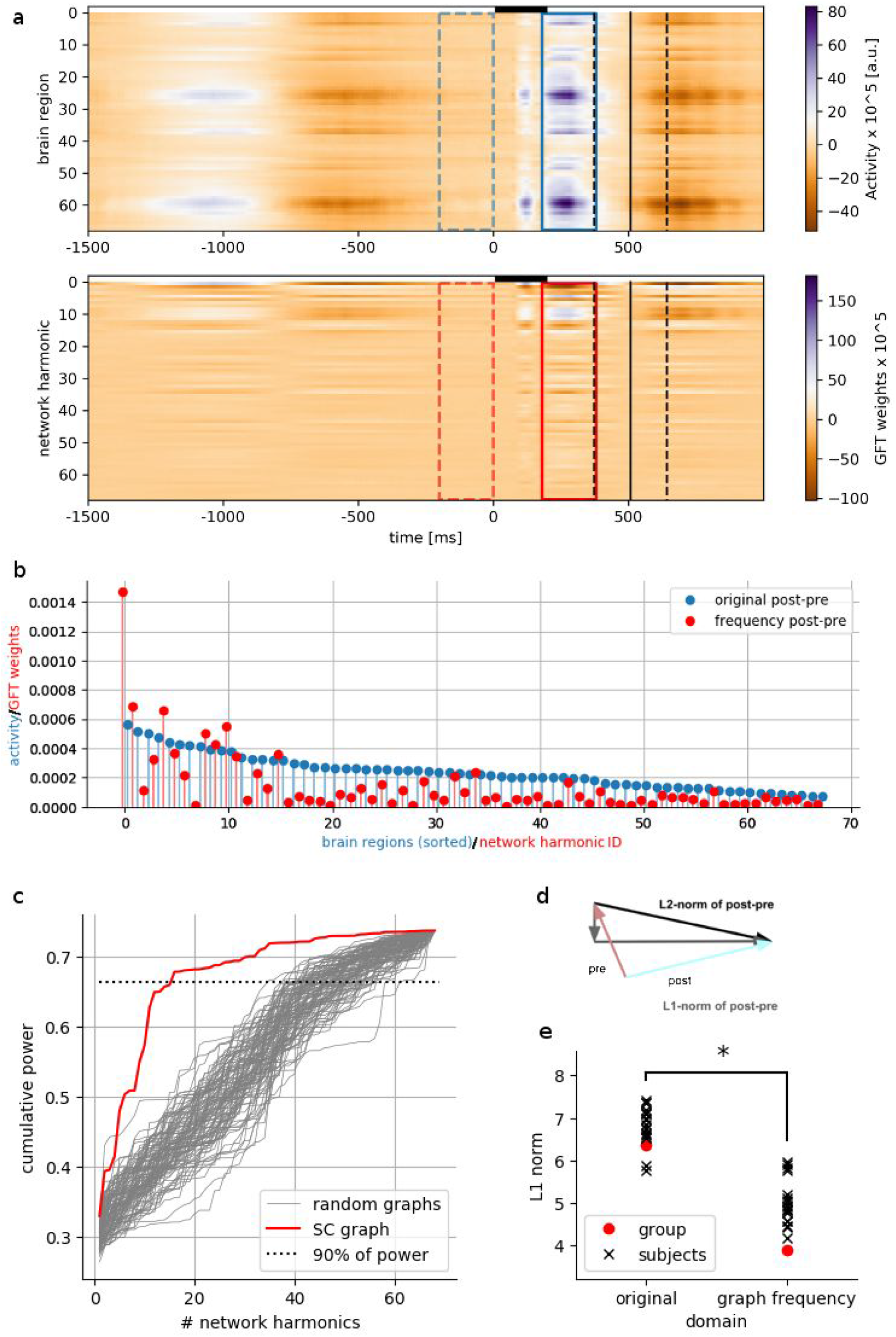
**a:** EEG signal averaged over all trials and all subjects in its original domain (top panel, rows are brain regions) and in the graph frequency/spectral domain (bottom panel, rows are network harmonics/eigenvalues). The windows which were used to assess the sparsity of the signal are marked as colored rectangles (dashed lines: pre-stimulus interval, solid lines: post-stimulus interval), as well as the interval in which the stimulus was presented (black bar, 0-200 ms) and the mean and standard deviation of the reaction times (solid and dashed black lines, respectively) across all trials and subjects. **b:** Signal averaged over the time points within the windows marked in panel a in the “original” and in the graph frequency domain. Brain regions are ordered by amplitude, network harmonics are ordered by eigenvalue. GFT: graph Fourier transform. **c**: Power captured cumulatively as more network harmonics are added, for the SC-derived graph (red line) and for 100 randomized graphs (grey lines). The dotted line marks 90% of the overall power. **d**: Illustration of how the L1 norm captures sparsity of the signal.The smaller the L1 norm, the more compact the signal, even as the L2 norm (power) remains the same. **e:** Difference between the L1 norms of the signals shown in panel b, both averaged over subjects, and, for illustration, for each subject. The star marks a significance difference between the means as assessed by a permutation test (see Methods).

We furthermore quantified the effect size of this difference. Since we used a nonparametric significance test, we did not use the usual method of Cohen’s d, as it relates to the t-statistic and assumes normal distribution of the underlying variables. Instead we employed the approach described in (Kraemer and Andrews 1982), which allows to compute a nonparametric effect size which is nonetheless interpretable in the same way as a t-statistic due to a transformation to normal distribution in the last step. The method is described in more detail in the next subsection (“*Statistical analysis of network harmonic activations (sliding window analysis)*”).

### 2.8. Statistical analysis of network harmonic activations (sliding window analysis)

For the results shown in Figures 3 and 4, we used a permutation test in order to identify network harmonics/brain regions that were more strongly activated (with positive or negative weight) during each post-stimulus window compared to a fixed pre-stimulus window. The post-stimulus windows covered the time course from 0 to 600 ms post stimulus onset, with a width of 50 ms (13 samples) and a 50% overlap between adjacent windows. We chose this approach in order to reduce noise and to control the number of multiple comparisons, while still preserving a good temporal resolution which allows direct assignment of significance to a certain time window and network/brain region.

**Figure 3.**
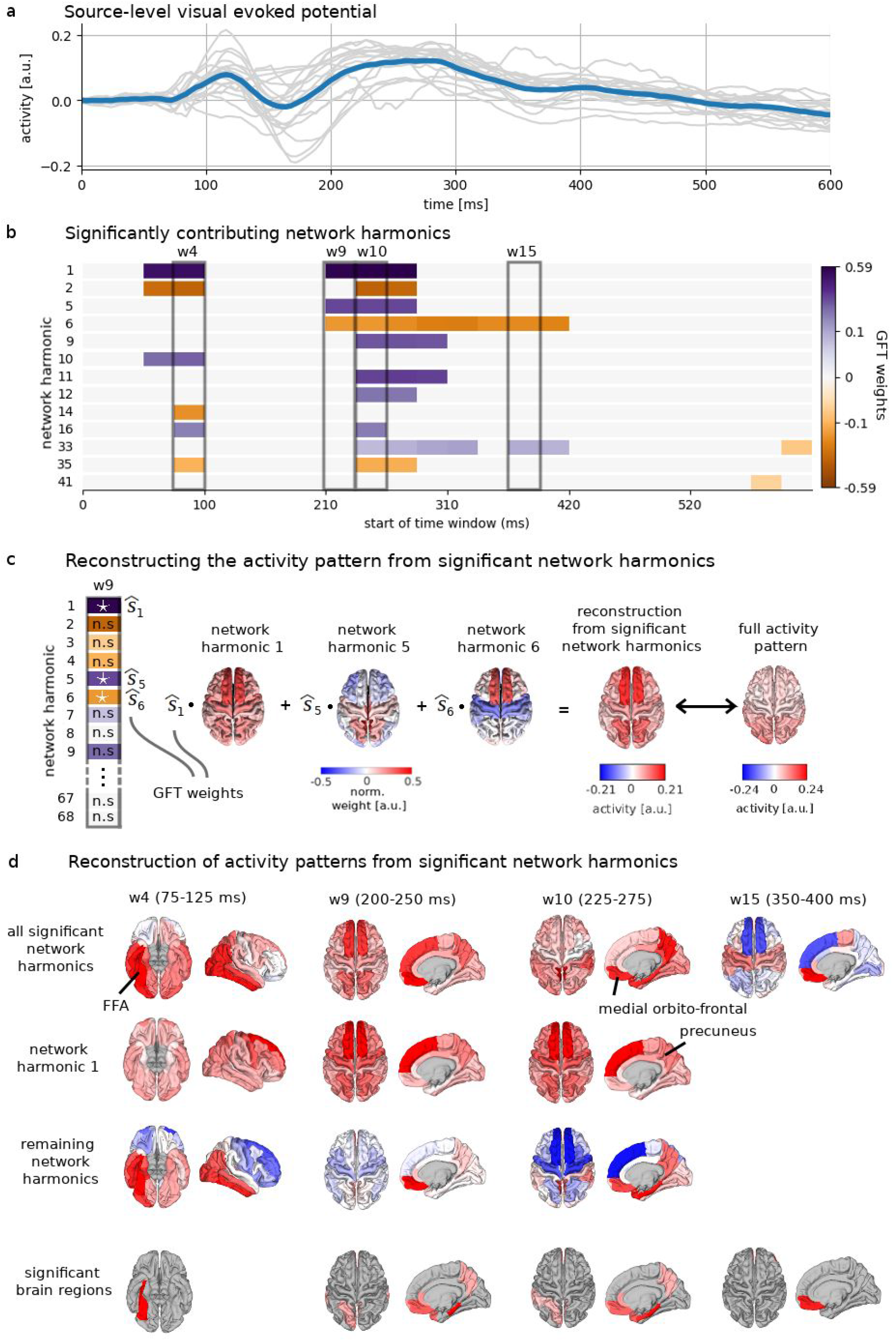
Network harmonic tracking during face detection task. **a:** EEG signal averaged over all brain regions and trials (event-related potential). Blue curve: average over all subjects. Gray curves: single subjects. **b:** Time courses of network harmonics for time windows in which they show a significant (de-)activation compared to pre-stimulus baseline. The gray boxes mark time windows (w) for which surface renderings are shown in panel **d**. **c**: Illustration of how activity (amplitude) patterns are reconstructed as a weighted sum (superposition) of those network harmonics that are identified as significant in the statistical analysis. The example of window 9 is shown: there is one GFT weight 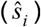 for each of the N=68 network harmonics *i*. GFT weights 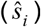 quantify the contribution of each network harmonic *i* to the activity pattern. The white stars mark those that are significant, the others are not significant (n.s.) and are not included in the weighted sum. The resulting reconstructed activity pattern is different from the full activity pattern because not all network harmonics are included. **d**: Surface renderings for four selected time windows. The activity patterns (amplitudes) reconstructed only from the significant networks (as illustrated in panel **c**) are shown in the top row. In cases where the first network harmonic was significant, this activity pattern is also shown as a superposition of this first network harmonic and the sum of all other significant networks (rows 2 and 3). The bottom row shows significant brain regions obtained with the same analysis done with the original signal (time courses as in panel **b** shown in Figure S3). FFA: fusiform face area. Precuneus and medial orbito-frontal regions are nodes of the default mode network (Raichle 2015).

**Figure 4.**
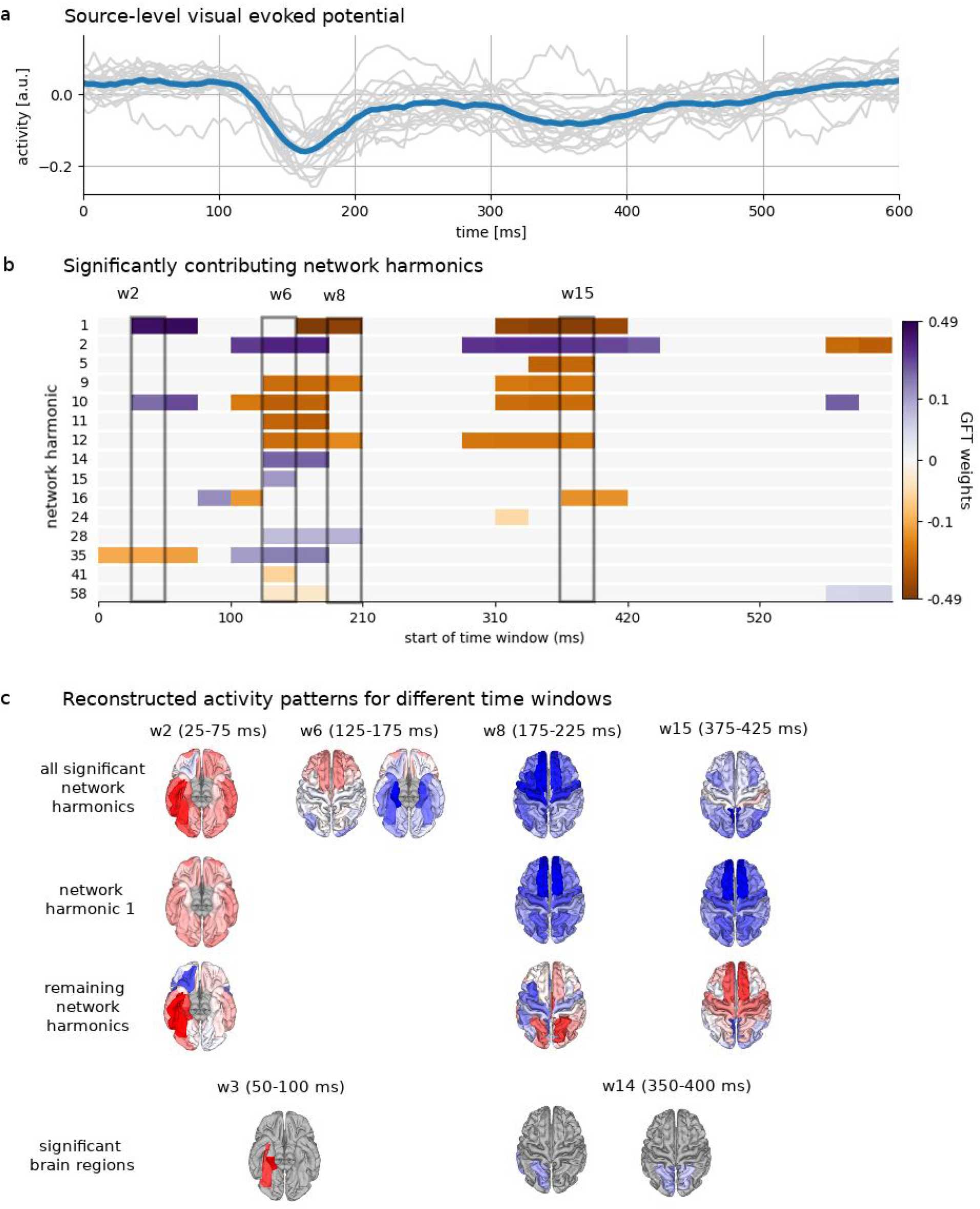
Differences between “face” and “scrambled” conditions. **a:** Difference between EEG signal averaged over all brain regions for each class of trials (event-related potential; “face” minus “scrambled”). Blue curve: average over all subjects. Gray curves: single subjects. **b:** Time courses of network harmonics for time windows in which they show a significant (de-)activation in “face” trials compared to “scrambled” trials. The gray boxes mark time windows (w) for which surface renderings are shown in panel **c**. **c**: Surface renderings for four selected time windows. The activity patterns reconstructed only from the significant networks are shown in the top row (see Figure 3c for an illustration). In cases where the first network harmonic was significant, this activity pattern is also shown as a superposition of this first network harmonic and the sum of all other significant networks (rows 2 and 3). The bottom row shows significant brain regions obtained with the same analysis done with the original signal. In two cases, the time windows selected are shifted by one, as there were not enough activations in the time window selected for the network harmonics (time courses as in panel **b** shown in Figure S4).

We computed individual differences for each post-stimulus window and then averaged across subjects, after normalizing each individual difference vector to length 1 in order to avoid the average being biased by subjects with higher signal power. The permutation consisted in switching, for a randomly selected subset of subjects, the pre- and post-stimulus intervals (i.e., the sign of the difference vector) and subsequently recomputing the average. The p-value was computed as the fraction of permuted averages showing a larger or equally large difference in activation as the empirical average. This was done on a single network/brain region level. We used a significance level of 0.05, corrected for multiple comparisons (N networks/brain regions and 23 sliding windows).

We also computed the nonparametric effect sizes for all time windows of the network harmonics that exhibited time windows of significant activation. This was done using the method described in (Kraemer and Andrews 1982) in the following manner for the comparison between pre- and post-stimulus intervals (illustrated in Figure S2A), and analogously for the comparison between faces and scrambled images:

1. Compute the difference in GFT weights between the (fixed) pre-stimulus interval and the selected post-stimulus sliding window. Obtain the L2-norm of the difference vector, and use it to normalize the pre- an post-stimulus GFT weights. This is necessary in order to make GFT weights comparable across subjects, and to make pre- and post-stimulus GFT weights comparable (left vs right panel of Figure S2A).
2. For a given network harmonic, select the median normalized pre-stimulus GFT weight. From the set of normalized pre-stimulus GFT weights, ordered along the subject dimension, select two more values to the left and right of the median, resulting in five “typical” normalized pre-stimulus GFT weights (red dots in Figure S2A, “pre”).
3. Determine the normalized post-stimulus GFT weights of the same subjects (red dots in Figure S2A, “post”).
4. Compute the median of these five normalized post-stimulus GFT weights. This can be seen as the “typical response” (blue dotted line in right panel of Figure S2A).
5. Determine the proportion of subjects whose normalized pre-stimulus GFT weights are smaller (“worse”) than the typical response (all subjects below the dotted line on the “pre” side in Figure S2A). This can be seen as the proportion of subjects whose post-stimulus activation of the chosen network harmonic was larger than their pre-stimulus activation; i.e. the proportional effect size. If the effect size is 1, this value is replaced by 1/(S+1), if it is S, it is replaced by S/(S+1) in order to avoid problems in the following step.
6. In order to obtain a measure that is readily interpretable in the same way as t-statistics, the nonparametric effect size is computed by evaluating the inverse cumulative standard normal distribution at the value of the proportional effect size. Note that if the effect is in the other direction, i.e. the GFT weight is significantly reduced, this will simply result in a negative effect size.

For the comparison between the two conditions (faces and non-faces/scrambled), replace “pre-” and “post-”interval with “faces” and “scrambled” intervals taken from the same post-stimulus sliding window. Also in this case, the normalization by the L2-norm of the difference vector is performed prior to computing the effect size.

Effect sizes are classified as for Cohen’s d, i.e. effect sizes up to 0.01 are very small, up to 0.20, small, up to 0.50, medium, up to 0.80, large, up to 1.20, very largen, and up to 2.0, huge.

## 3. Results

### 3.1. Network harmonics: Building blocks of brain activity derived from structural connectivity

We use a multimodal approach to large-scale spatiotemporal dynamics, building on the finding that functional connectivity (FC) is partly explained by the structure of anatomical long-range connectivity (structural connectivity, SC) in the brain, both in fMRI and M/EEG (Vincent et al. 2007; Hagmann et al. 2008; Honey et al. 2009; Damoiseaux and Greicius 2009; Deco et al. 2013; Cabral et al. 2014; Goñi et al. 2014; Tewarie et al. 2014; Atasoy et al. 2016; Meier et al. 2016; Glomb et al. 2017; Abdelnour et al. 2018; Tewarie et al. 2019; Chu et al. 2015; Wirsich et al. 2017). The key is the mathematical framework of graph signal processing (GSP) which provides mathematical tools that allow to extract harmonic basis functions from the SC, which then serve to obtain a graph-spectral representation of the data (Shuman et al. 2012). We term these basis functions network harmonics. The technique that we use to extract network harmonics is illustrated in Figure 1a. The first step is to compute a consensus (average) SC matrix (see Methods), which encodes a graph whose nodes are the brain regions and whose edges are white-matter connections (illustrated below the SC matrix in Figure 1a). Next, the graph Laplacian is obtained from the SC matrix in one simple step; it encodes the same information as in SC. Finally, the eigenvectors of the graph Laplacian are obtained. These eigenvectors are the basis functions of the SC graph, i.e., the network harmonics. Figure 1b shows the first four network harmonics, corresponding to the smallest four eigenvalues. The first ten network harmonics are shown in Figure S1. For illustrative purposes, we show the network harmonics using three equivalent representations: as a color-coded eigenvector, where each element corresponds to a brain region; projected onto the brain surface, i.e. the elements of the vector plotted in anatomical space; and as node weights using the same graph shown in Figure 1a.

In these representations, nodes that have a similar color are nodes that are close (i.e., similar) in “connectivity space”: they share patterns of connectivity to the network as a whole, on a certain graph frequency. Note that this graph frequency is a spatial frequency, unlike in the most common case of obtaining the Fourier spectrum of a temporal signal In the remainder of the paper, we will use the term “graph frequency” to clearly distinguish it from the more common use of temporal frequencies. The number of graph frequency bins is determined by the number of brain regions in the parcellation (here, N=68).

Starting from the eigendecomposition, the underlying graph can be interpreted as a superposition of rank-1 matrices obtained by computing the outer product of each eigenvector with itself, weighted by its corresponding eigenvalue (Figure 1c). Large positive values in this outer product (which is an *N* by *N* - matrix just like the Laplacian/SC itself) only occur where two entries of the eigenvector which both have large weights of the same sign are multiplied together. Thus, two brain regions which are close together on the eigenvector encode a strong connection. We illustrate this point further by computing the correlation between the superposition of outer products - the reconstruction - and the original graph Laplacian (see Methods for details). Figure 1d shows that adding low-frequency network harmonics results in a negative correlation between these matrices, −0.75 being the minimum, reached when using the first 21 network harmonics. Note that positive weights corresponding to connected regions result in a negative correlation to the (all-negative) Laplacian edge weights. Thus, the first 21 low graph frequency network harmonics are most important for capturing integration in the SC-derived brain network. On the other hand, the correlation jumps from 0.03 when using 61 network harmonics to 1 when all network harmonics are used, indicating that the network harmonics with the highest graph frequencies meaningfully capture segregation (positive correlation to the graph Laplacian).

The first network harmonic, shown in Figure 1b, possesses, per definition, the lowest graph frequency. The difference between pairs of nodes which are connected according to the SC matrix is minimized. Formally, the sum of differences between connected pairs is quantified by the *smoothness* of each eigenvector (see Methods); thus, the ordering of network harmonics by graph frequency results from their ordering by smoothness. Indeed, the first network harmonic is highly correlated with the strength (sum of edge weights) of each node (r=0.94,p=6*10^−33^). The second one describes a connectivity gradient along the frontal to occipital axis of the brain; the third separates the two hemispheres; the forth network harmonic divides the parietal lobe from the rest of the brain. As the eigenvalue increases, the graph frequency also increases, resulting in finer-grained subdivisions of the cortex. Smoothness is thought to be meaningful because evolutionarily, it makes sense that connected regions should also be functionally related and should thus co-activate (Gu et al. 2018; Tomasi, Wang, and Volkow 2013). It is important to clarify that while some of the network harmonics are easily interpretable, as in the examples given above, this is not always the case as the SC-derived brain graph is complex. Instead, network harmonics should be understood as analytically obtained building blocks which function cumulatively, as illustrated above with the correlation between reconstructed and original connectivity matrices.

Figure 1e illustrates that, as a result of this multi-scale network representation, two nodes can be very similar in one network harmonic, and very dissimilar in another, indicating that on one graph frequency, the two regions have similar connectivity patterns, while on another, they have dissimilar patterns. This reflects the fact that each brain region fulfills multiple functions which can only be understood in conjunction with inputs and outputs from other brain regions, i.e., a functional network (Fox et al. 2005).

Taken together, the network harmonics used here capture the multi-scale and hierarchical properties of brain networks (Betzel and Bassett 2017; Yeo et al. 2011).

### 3.2 Network harmonics derived from structural connectivity provide a sparser basis for the EEG signal

As with sine and cosine waves of different frequencies in the well-known Fourier transform used for one-dimensional continuous time series, network harmonics form an orthonormal basis in which any signal can be approximated to arbitrary precision, provided it is defined on the same graph from which the network harmonics were extracted. We use source-projected EEG task data from a visual tasks which involves recognizing images of faces (see Methods). In this view, network harmonics provide building blocks of the cortical activation patterns observed during task performance. “Activation pattern” here refers to a set of all N=68 brain region’s signal amplitudes - or activities - measured at a certain point t in time (amplitudes are obtained by projecting EEG activity from the scalp into the gray matter [see Methods for details]). Using the network harmonics in this way, the EEG signal is transformed into the orthonormal function basis given by the network harmonics via the graph Fourier transform (GFT, Shuman et al. 2012; Figure 2a). This results in a spectral representation of the activation patterns in terms of GFT weights which quantify how much each network harmonic contributes to the activation pattern. One such spectral representation is obtained for each point in time. In the inverse direction, each activation pattern can be reconstructed as a weighted sum of the network harmonics, without loss of information.

In Figure 2b, the average signal amplitude of each brain region is shown in blue for the time window marked with solid lines in Figure 2a, after the pre-stimulus activation pattern has been removed (marked by the dashed lines in Figure 2a). We have ordered the brain regions by amplitude in order to show the comparison with the GFT weights of the spectral representation, which are shown in red and which are naturally ordered by their eigenvalue. The distribution of activity is different from that of GFT weights (note that the overall power - L2-norm - is the same, which allows us to directly compare the two domains in this way). While in the original signal, activity is distributed throughout the entire cortex, in the domain of the network harmonics, the GFT weights fall off quickly as the eigenvalue increases: the first 10 network harmonics capture 57% of the power, and the first 15 network harmonics are sufficient to capture 90% of the power (red line in Figure 2c). In other words, most of the activity is captured by the first few - smoothest - network harmonics.

We compare this capacity of the SC graph to capture important statistical dependencies in the EEG signal to that of 100 randomized graphs, in which the sequence of node strengths (sum over all edges connected to a node) is preserved. Since the node identities are the same, and represent brain regions in the same way as the SC-derived graph, the EEG signal can indeed be expressed perfectly by the random network harmonics extracted from a given random graph if all N=68 network harmonics are used; however, since the underlying graph does not reflect the true structure of the “brain graph”, we expect this representation to be less efficient. We find that in all of the randomized graphs, the number of network harmonics (ordered by eigenvalues) that is necessary to capture 90% of the power of the signal exceeds that of the SC graph (Figure 2c, grey lines), with a median number of necessary network harmonics of 46 (minimum: 34, maximum: 60) as opposed to 15 when the real, SC-derived network harmonics are used. Next, we directly test whether the graph frequency domain provides a quantifiable advantage over the original signal domain. It is not possible to compare the two domains in terms of how many dimensions (i.e., number of brain regions in the case of the “original” domain, number of network harmonics in the case of the graph frequency domain) are necessary to capture a certain amount of power of the signal, as it would be unclear which brain regions should be removed from the activation patterns, and how that should be interpreted. Instead, we employ the concept of sparsity, which is used in the framework of compressed sensing (Candès et al. 2006; Candes and Romberg 2006). Sparsity quantifies the intuitive idea that a “good” function basis allows the signal to be captured by a low number of dimensions. In practice, this is achieved by optimizing the L1-norm of the signal representation, i.e. the sum of absolute values in the signal (Ramirez et al. 2013). Note that the power of the signals (L2-norm) is the same, which makes this a valid comparison, as no information is lost when the signal is transformed between the two domains. Figure 2d illustrates how L1-norm is used to quantify sparsity. We compute the L1-norm of the signals as shown in Figure 2b, which represent the difference between pre- and post-stimulus intervals (windows in Figure 2a) in both signal domains. The L1-norm of the network harmonics-based difference vector is significantly smaller than the L1-norm of the difference vector of the original signal (Figure 2e, mean distances significantly different on a level of α=0.05 [two-sided test], Monte Carlo simulations with 10.000 permutations). Computing the nonparametric effect size corresponding to this difference indicates a very large effect (effect size of 1.6, see Methods for details). This is consistent with the idea that the difference between the two intervals is captured by fewer dimensions in the network harmonics-representation than is the case in the original representation.

This suggests that the ordering by smoothness captures important features of the functional signal; that the least smooth networks may be representing predominantly noise; and that thus, the domain of harmonic networks could be used for dimensionality reduction and filtering.

### 3.3. Network harmonics are able to track EEG task activity

We next investigate whether network harmonics are a functionally meaningful basis that is able to explain spatiotemporal dynamics of brain activity recorded during a facial recognition task. Note that the fact that network harmonics form an orthonormal basis means that any signal on the graph, and hence any EEG activation pattern, can be perfectly captured. Our question is therefore, first, whether a small number of network harmonics suffices, because this would mean that the SC-derived network harmonics are related to functional networks/co-activation patterns present in the functional data (Abdelnour et al. 2018; Meier et al. 2016; Tewarie et al. 2014) Second, whether this approach provides any advantage over the more conventional approach of testing each brain region one by one for significance.

We transform EEG data into the space spanned by the network harmonics for each point in time from the average over trials of each subject (Figure 2a). This results in N time courses of GFT weights, one for each network harmonic (Figure 2a). Next, we identify those network harmonics that significantly contribute to the processing that takes place in response to the visual stimulus by applying a permutation test (see Methods) that compares the pre-stimulus to post-stimulus GFT weights averaged over 50 ms sliding windows (50% overlap, see Methods for details). Figure 3a and b show the source-space visual evoked potentials (signals averaged over all brain regions; VEPs) and the time courses of the network harmonics with significantly different GFT weights, respectively. Only 13 out of 68 network harmonics contribute significantly (Figure 3b). Figure S2B shows that the nonparametric effect sizes for all the network harmonics and time windows which were identified as significant are huge (>1.2) or very large (>0.8) with the exception of network harmonic 33 in the very last time window, where the effect size was large (>0.5; see Methods).

Next, we use only the significant GFT weights to reconstruct cortical activation patterns from which non-significant distributed patterns of activity have been removed, as illustrated in Figure 3c; i.e., we compute the sum of only the significant network harmonics, weighted by their GFT weights (see Methods, equation (6)). This resulting reconstructed signal is thus again in the “original” domain, i.e. it is an activation pattern consisting of amplitudes (“activity”). Figure 3d shows surface renderings of activation patterns that were reconstructed using only the significant network harmonics (by applying the inverse GFT). There are two intervals that exhibit significant network harmonics, an early one (~50-125 ms after stimulus onset) and a later one (~210-430 ms). During the early time interval, there is a general increase in activation (amplitude), especially in the occipital cortex. The right fusiform area (FFA) is visible, consistent with the finding that face processing is lateralized to the right hemisphere (Meng et al. 2012; Rossion, Schiltz, and Crommelinck 2003). The later time interval exhibits initially a strong activation in prefrontal areas following the end of the stimulus presentation at 200 ms, and later, starting at around 300 ms, activity in the somatosensory/motor areas corresponding to preparation of the motor response.

One advantage of using the network harmonics in this analysis is that the observed activation pattern can be understood as a superposition of network harmonics. Specifically, it is useful to remove the contribution of the first, smoothest network harmonic which results from a non-specific increase in cortical activation (Figure 3d, second row). This way, the activation of visual areas in the occipital cortex in general, and of the right FFA in particular, in the early time interval is even more prominent (Figure 3d, w4, third row). In the later time interval, an increase in GFT weight of the first network harmonic is present up to around 300 ms. When discounting this increase, deactivation of superior frontal regions starting at ~230 ms, and activation of precuneus and medial orbito-frontal regions - both prominent nodes of the default mode network (Raichle 2015) - starting at 210 ms is revealed (Figure 3d, w9 and w10, third row).

We conduct the same analysis in the original signal domain, i.e. using brain regions instead of network harmonics. The bottom row in Figure 3d shows a series of surface renderings that mark the brain regions whose amplitudes are found to be significantly different from the pre-stimulus interval within the analyzed time windows (see Figure S3 for all significant time courses). While in the early time interval, the activation of the FFA is present as observed in the activity patterns resulting from reconstruction from significant network harmonics, this is not the case for the activation of frontal regions nor for the activity in somatosensory/motor regions around 200 ms.

### 3.4. Network harmonics for distinguishing between faces and non-faces

So far, we have shown results for the processing of images of faces. In the same experiment, roughly the same number of trials involved showing scrambled versions of these images (see Methods for details). We used network harmonics to identify differences between these two conditions, which is a way to determine which components of the observed spatiotemporal dynamics are specific to images of faces, as opposed to visual stimuli in general. Thus, we compute the difference between EEG activity found during “face” trials and subtract activity from “scrambled” trials. Figure 4a and b show the VEPs and the significant network harmonics’ time courses, respectively.

We identify three intervals which exhibit significantly different GFT weights between the two trial types and reconstruct activity patterns (amplitudes) by transforming back into the “original” domain as above.

In the earliest interval (~25-100 ms; note that the time window starting at 25 ms extends up to 75 ms), we observe a positive amplitude pattern in the reconstructed cortical activity (Figure 4c, w2, first to third row), indicating stronger overall activation of almost the entire cortex in “face” trials as opposed to “scrambled” trials. Removing the first network harmonic’s contribution as above, we find that the remaining contributing network harmonics superimpose to exhibit a strong activation specifically of the right FFA in “face” trials (Figure 4c, w2, third row). We also compute the nonparametric effect size (see Methods for details) and find that the effect size is huge (>1.2) for network harmonic 10 (see Figure S1 for a surface rendering of this network harmonic), while it is very large (>0.8) for all other involved network harmonics except network harmonic 35, for which the effect is only large (>0.5) to medium (>0.2).

In the middle interval (~100-230 ms), the polarity of superior frontal regions reverses from positive (stronger activation in “face” trials) to negative (stronger activation in “scrambled” trials) over the course of the 4 time windows that make up this interval (Figure 4c, w6 and w8, first row). Looking at the first network harmonic, which only contributes during the second half of this interval (w8), it becomes clear that an overall decrease of activation contributes to this observation. In this interval, the effect size is huge for network harmonics 2, 10, 11, 12, 15, 28, and 41 for the time window extending from 125 to 175 ms, and even longer for network harmonics 10, 11, 28, and 41 specifically. During these time windows, negative activation is observed in the FFA (Figure 4c, w6). This corresponds to the well-known N170, a strong negative inflection which is robustly observed when images of faces are presented (Bentin et al. 1996; Eimer 2000). Effects are further very large (>0.8) or large (>0.5) for network harmonics 1, 9, and 58. Only small or very small effects are observed for the earlier time window of this interval (100-150 ms).

In the late interval (~280-460 ms), there is a pattern that is consistent with overall less activity in “face” trials as opposed to “scrambled” trials, with this deactivation becoming stronger in occipital areas over the course of the interval (Figure 4c, w15, first row). When removing again the contribution of the first network harmonic, it is revealed that starting at ~340 ms, there is stronger activity in frontal regions in “face” trials, as well as in somatosensory/motor regions (Figure 4c, w15, third row), indicating there may be a difference in response behavior between the conditions. In terms of effect sizes, network harmonics 1, 2, 5, 9, 10, 12, and 16 exhibit huge effect sizes (>1.2), most extensively for network harmonics 2, 5, 9, and 10. Most other network harmonics whose activation is significantly different between “faces” and “scrambled” trials in this interval have a very large effect size (>0.8), with the exception of the early time window (starting at 310 ms), for which effect sizes are very small (>0.01) to medium (>0.2).

As before, analysis of the original signal (using the brain regions themselves) does reveal stronger activation of the FFA in “face” trials in the early time interval, but in the later two intervals, none of the frontal or somatosensory/motor differences are present.

## 4. Discussion

Spatiotemporal dynamics of large-scale networks in the brain have been shown to be highly relevant both in the healthy human brain as well in many brain disorders. EEG is a powerful tool for mapping and understanding fast network dynamics in the human brain, as it does not only record direct brain activity on a sub-millisecond time scale, but is also comparatively cheap as well as portable, and thus, at least in theory, well-suited for clinical applications. However, EEG suffers from the effects of volume conduction, which makes it difficult to compute functional connectivity and derive valid functional networks from it.

We introduce network harmonics - Fourier basis functions of the brain structural connectivity graph - as building blocks of EEG source-level activity. We leverage well-understood tools from graph signal processing, and the theory of harmonic modes, to map fast, large-scale cortical dynamics in source-projected EEG data. In particular, graph signal processing allows us to obtain meaningful “building blocks” of functional activity from the graph of structural connectivity without the need to compute functional connectivity.

We explicitly show that network harmonics meaningfully capture integration and segregation in the brain network graph, and demonstrate their efficiency compared to the ROI-by-ROI-representation of the EEG signal. We conduct a statistical analysis of EEG data recorded during a facial recognition test, which we perform in the domain of network harmonics. We show that a few network harmonics are sufficient to capture task dynamics.

### 4.1. Combining data modalities to probe the structure-function relationship in brain networks

Using network harmonics in the way described here combines data from multiple modalities, namely we decompose functional (EEG) data using building blocks derived from structural connectivity (diffusion MRI). Our main finding is that EEG functional activity can be efficiently expressed as a superposition of co-activation patterns directly extracted from structural connectivity, adding to the evidence which suggests that function is shaped by structure in the brain on a macroscopic scale, and confirming that this finding extends to EEG (Chu et al. 2015; Finger et al. 2016; Wirsich et al. 2017; Glomb, Mullier, et al. 2019).

Our approach also offers a number of practical advantages. On the one hand, it is unnecessary to compute functional connectivity, which is impacted by volume conduction, i.e., the spread of electric fields through brain tissue that results in spurious statistical relationships between time series recorded from different brain regions. On the other hand, it allows us to take a statistical approach which does not rely on making any prior assumptions on where in the brain differences are expected. Instead, we use information on integration and segregation between brain regions as encoded in the structural connectivity matrix.This also means that we identify significant patterns of co-activation involving the entire cortex, while in an approach where each ROI is treated independently, the activity in all non-significant brain regions is essentially ignored. This is in line with the idea that cognitive functions are fulfilled not by single regions, but by networks (Fox et al. 2005). In a similar vein, even though in many applications, the cortex is partitioned into a fixed number of non-overlapping networks (Yeo et al. 2011), it has long been appreciated that the brain network, both functional and structural, consists of overlapping hierarchical modules (Betzel and Bassett 2017). Thus, we take advantage of the multi-scale, hierarchical network structure encoded in the SC matrix in order to decompose and analyze EEG functional data.

It is important to mention that these building blocks are not conceptually equivalent to “brain states”. Instead, a combination of network harmonics is assumed to contribute simultaneously at any given point in time, and the degree to which each network harmonic contributes is allowed to vary on a continuous scale. In this sense, our approach does not assume abrupt changes between states, in contrast to, for example, microstates (Van de Ville, Britz, and Michel 2010), or Hidden Markov Model-based approaches (Baker et al. 2014; Quinn et al. 2018), where each point in time is assigned to exactly one state.

Furthermore, the relationship between structure and function in the human brain is one of the major topics of contemporary neuroscience which is also pursued in the field of connectomics. Converging evidence suggests that functional connectivity is in part determined by structural connectivity, one of the most robust findings being that brain regions which have a direct anatomical link in the SC have stronger FC than those that do not (Chu et al. 2015; Finger et al. 2016; Glomb, Mullier, et al. 2019). Furthermore, generative models often use SC as an underlying scaffold that shapes the simulated functional activity (Deco et al. 2013; Honey et al. 2009; Messé et al. 2015; Wang et al. 2019). Our approach could easily be used, for example, to explore the impact of alterations in the SC on the resulting network harmonics and their relationship to simulated FC. Beyond these basic findings, harmonic modes of the SC have been shown to explain patterns of activity-dependent disease propagation (Raj et al. 2020), and recent work done with fMRI suggests that the degree to which functional activity is aligned with harmonic modes of the SC is relevant for cognitive flexibility (Medaglia et al. 2018) and functional specialization (Preti and Van De Ville 2019). Note that the lower spatial resolution and SNR of EEG, compared with fMRI, make these results only partially applicable here. Specifically, it is likely that network harmonics with a high graph frequency are not meaningful for the functional data, as local activity is always highly dependent on the activity of neighboring regions. This is in line with our finding that high graph frequency networks contribute very little to the task activity.

As an outlook, combining several data modalities is an approach that is becoming more and more popular in the field of personalized medicine, where large amounts of data are compiled for the same individual for the purpose of diagnosis, treatment planning, and prognosis via machine learning techniques. While the present study analyzes data on the group level, our approach holds the potential to be developed for the individual level by using information from individual connectomes (here we used a consensus connectome from a different group of subjects), as well as including microstructural information from structural scans.

### 4.2. Harmonic modes provide a powerful theoretical framework

On the methodological side, network harmonics are firmly rooted in the theoretical framework of harmonic modes, linking them to Fourier basis functions in other domains such as sine and cosine waves on the circle and spherical harmonics on the sphere (Robinson et al. 2016; Gabay, Babaie-Janvier, and Robinson 2018). A basic property of harmonic modes is that they are ordered by smoothness, and in the context of neural data, this means that they provide a multi-scale representation of the brain, reflecting the hierarchical organization of brain networks (Margulies et al. 2016; Betzel and Bassett 2017; Glomb, Kringelbach, et al. 2019).

Graph signal processing provides well-developed tools to study signals using harmonic modes of graph domains (Shuman et al. 2012). This allows for our method to be simple and fast. Here we have only used the most basic of these tools, and more advanced possibilities remain to be explored, in particular, designing and applying filters in the spatial spectral domain in order to remove noise and to explore the relationship between temporal and spatial frequencies in a joint spectral representation.

Beyond the link to basis function sets in other domains (circle, sphere), eigenvectors of the Laplacian are encountered when solving differential equations that link space and time in continuous domains, for example, the wave equation or the diffusion equation. In this theoretical context, the eigenvectors of the Laplacian are the spatial solutions; for example, in vibrating systems, they constitute standing wave patterns. There is also a direct theoretical link between the normalized graph Laplacian and Markov chains, i.e. a model of temporal evolution. The link to the temporal domain is established via the wave speed, i.e., the speed with which a wave propagates in space. In the brain, the link between spatial and temporal frequencies is conceivably established by the delays between brain regions derived from conduction speeds and fiber lengths/geodesic distances, which have been shown to crucially shape neural activity (Cabral et al. 2014). This way, the network harmonics shown here have a theoretical link to temporal frequencies, even though we do not explicitly consider conduction speeds. Although conduction speeds cannot be assumed to be uniform across the brain, in theory, each network harmonic could be linked to a (range of) temporal frequencies. Recent work has shown that oscillations, which play a major role in EEG analysis and in the functioning of the brain in general (Fries 2015; Klimesch 1996; Başar et al. 2000) might in part be explained by harmonic modes of the SC (Raj et al. 2020). (Glomb, Kringelbach, et al. 2019; Raj, Kuceyeski, and Weiner 2012; Gabay, Babaie-Janvier, and Robinson 2018). An appropriate model could enable estimation of these speeds. Research in this direction is just beginning to emerge (Gabay, Babaie-Janvier, and Robinson 2018; Atasoy et al. 2017; Atasoy, Donnelly, and Pearson 2016; Raj et al. 2020).

### 4.3. Limitations

The main limitation of network harmonics is that the structural connectivity matrix contains many false-positives as well as false-negatives, the latter especially when it comes to long-range and cross-hemispheric connections. This is also the reason why we used a consensus SC matrix instead of individual connectomes - taking into account information from the whole population makes the connections more reliable. At the same time, even if diffusion MRI and tractography algorithms were able to correctly identify all white matter connections in an individual brain, the fiber counts obtained in this manner do not (necessarily) correspond to the effective impact that one brain region has over another; likewise, connections in the SC are undirected. Therefore, the network harmonics used here cannot be seen as canonical at this point, as it is still unclear in how far they depend on the exact methodology and quality of the SC graph used. Furthermore, while we do not compute functional connectivity and thereby “sidestep” the issue of volume conduction to some degree, its impact is of course not removed from the functional data, and spurious correlations between nearby brain regions are still going to be reflected in the combinations of network harmonics that are found for each point in time. This also reveals a more general problem, regarding the absence of ground truth, which makes it difficult to validate any method that aims at tracking network activity. Showing relations to behavioral measures, including alterations in patient populations, would be useful here and should be addressed by future work.

Lastly, the nodes of the network harmonics are defined based on a standardized parcellation (Desikan et al. 2006). It is unclear whether the size, number and shape of the brain regions defined in this atlas are optimal for EEG source analysis in general (Farahibozorg, Henson, and Hauk 2018), and for network harmonics in particular. Future work could test whether optimization of the parcellation towards higher sensitivity and specificity to the signal of interest is possible.

### 4.4. Conclusions and future work

In this study, we show how network harmonics establish a link between connectomics and a rich and general mathematical theory. We demonstrate that graph signal processing is a methodological framework which is easy to use, statistically powerful, and holds a large potential for exploring the relationship between structure and function specifically in the context of EEG. Future work is necessary in order to understand how basis functions derived from structural or functional connectivity relate to each other, but also to the cortical microstructure, to behavioral measures, and to alterations in disorders. On the other hand, applying the framework described here to a multitude of tasks is straightforward and should produce new insights into large-scale dynamics as measured with source-projected high-density EEG.

Of particular interest is the relationship between temporal and spatial frequencies, for which EEG data with their high temporal resolution should prove particularly valuable. Graph signal processing provides the tools and theory necessary to pursue this direction, and should be linked to modelling work being done in this direction.

## Acknowledgements

This work was supported by Swiss National Science Foundation Sinergia grant no. 170873 and grant no. 169198 (to S. Vulliemoz), PP00P1_183714 to G. Plomp.

## Notes

### Competing Interest Statement

The authors have declared no competing interest.

