## Supplementary Information for "Connectome spectral analysis to track EEG task dynamics on a subsecond scale"

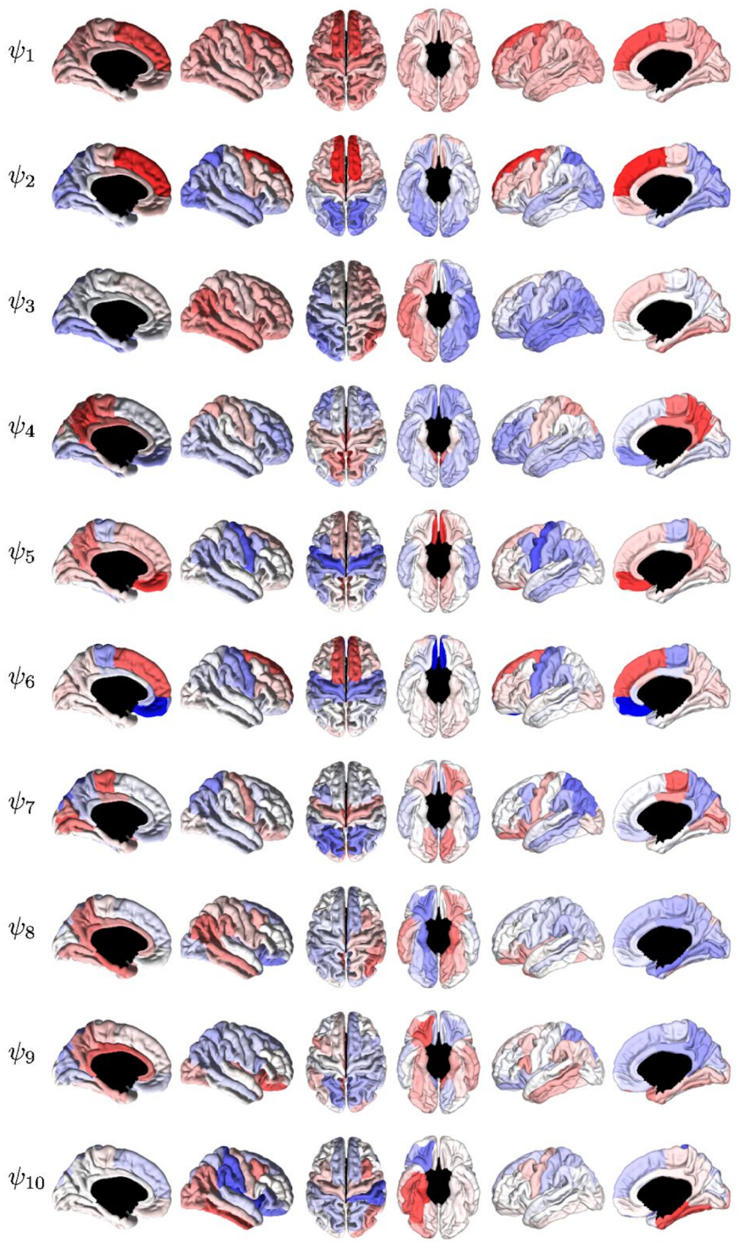


Figure S1: The first 10 harmonic networks plotted on the brain surface, viewed from different angles.

**A** - Illustration of statistical approach

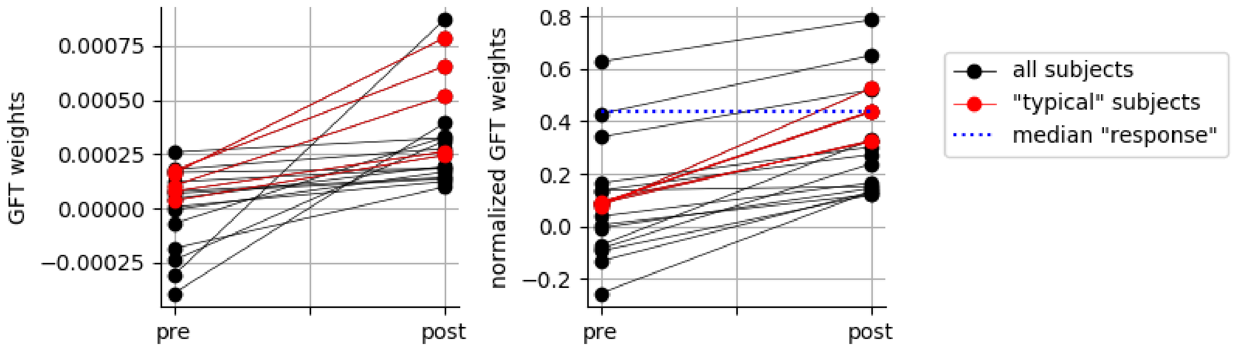


**B** - Nonparametric effect sizes: pre- vs. post-stimulus interval

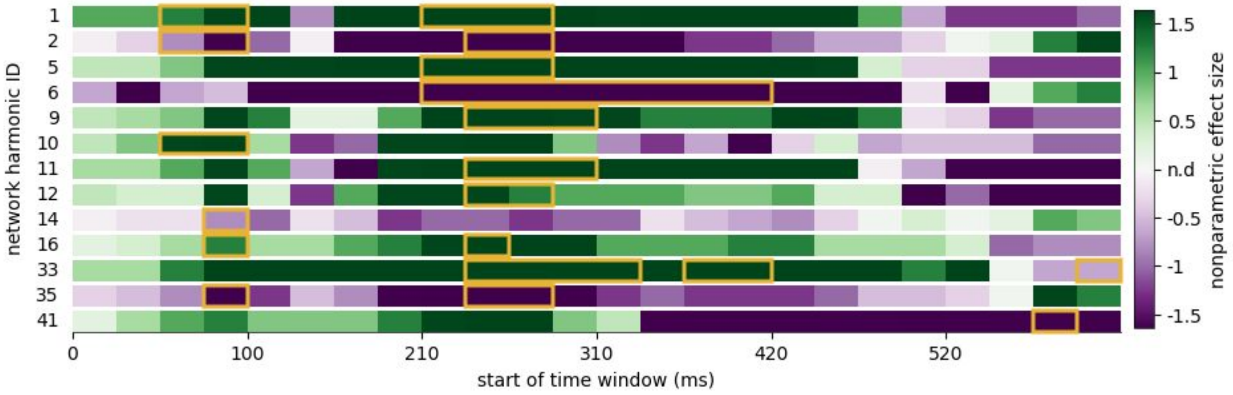


**C** - Nonparametric effect sizes: faces vs. scrambled trials interval

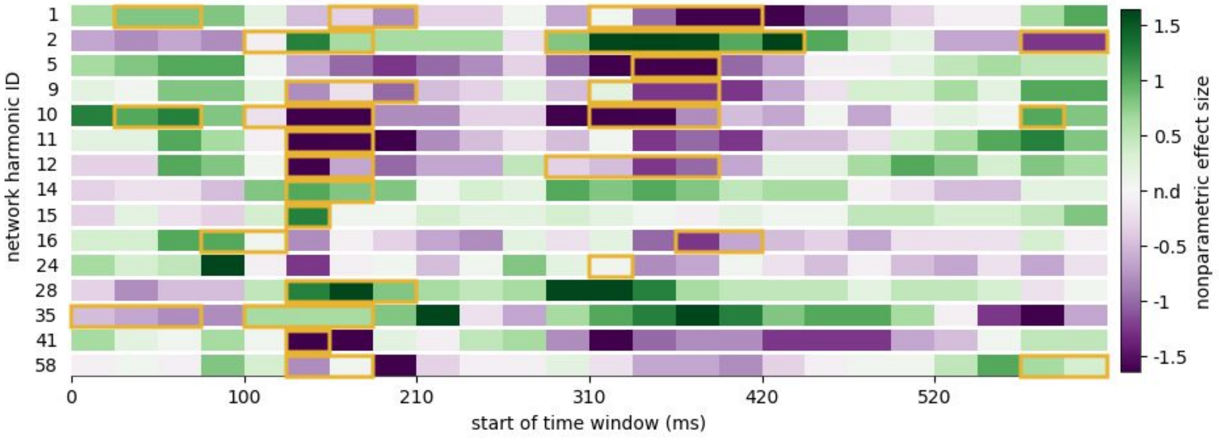


Figure S2: Nonparametric effect sizes for differences in GFT weights (activation of network harmonics) for network harmonics which exhibit time windows of significant differences (marked in yellow). **A** - Illustration of nonparametric effect size (Kraemer and Andrews 1982) applied to the comparison between pre- and post-stimulus activations of network harmonics. Left panel: Without normalization of GFT weights by the norm of the difference between pre and post. Right panel: With normalization. **B** - Nonparametric effect sizes for GFT weight differences between pre- and post-stimulus interval, corresponding to the activations shown in Figure 3. **C** - Nonparametric effect sizes for GFT weight differences between faces and scrambled faces trials, corresponding to the activations shown in Figure 4.

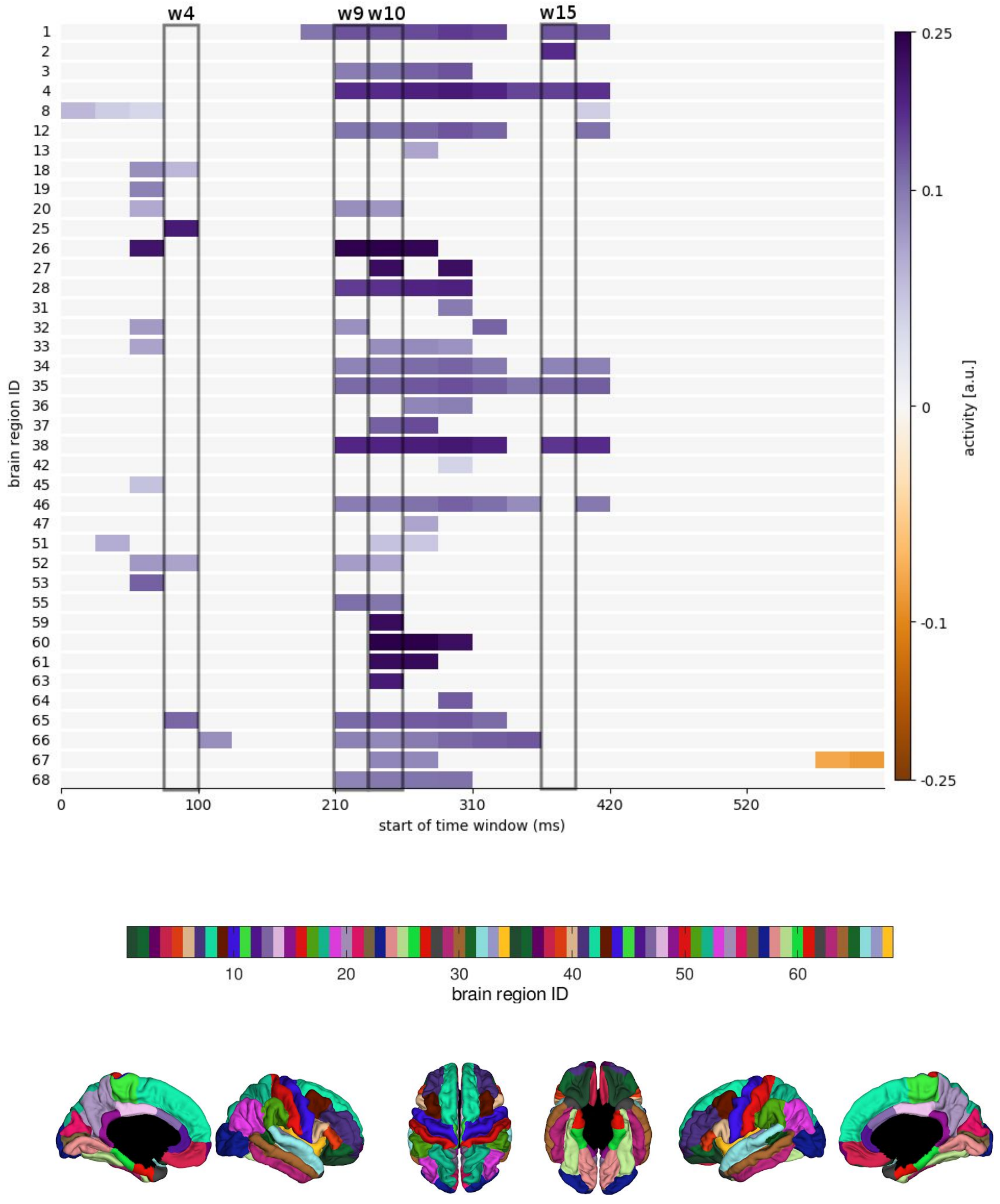


Figure S3. (related to Figure 3) Time courses of significantly contributing brain regions. Gray boxes mark time windows for which surface renderings are shown in Figure 3. Brain regions are color-coded in the plot at the bottom.

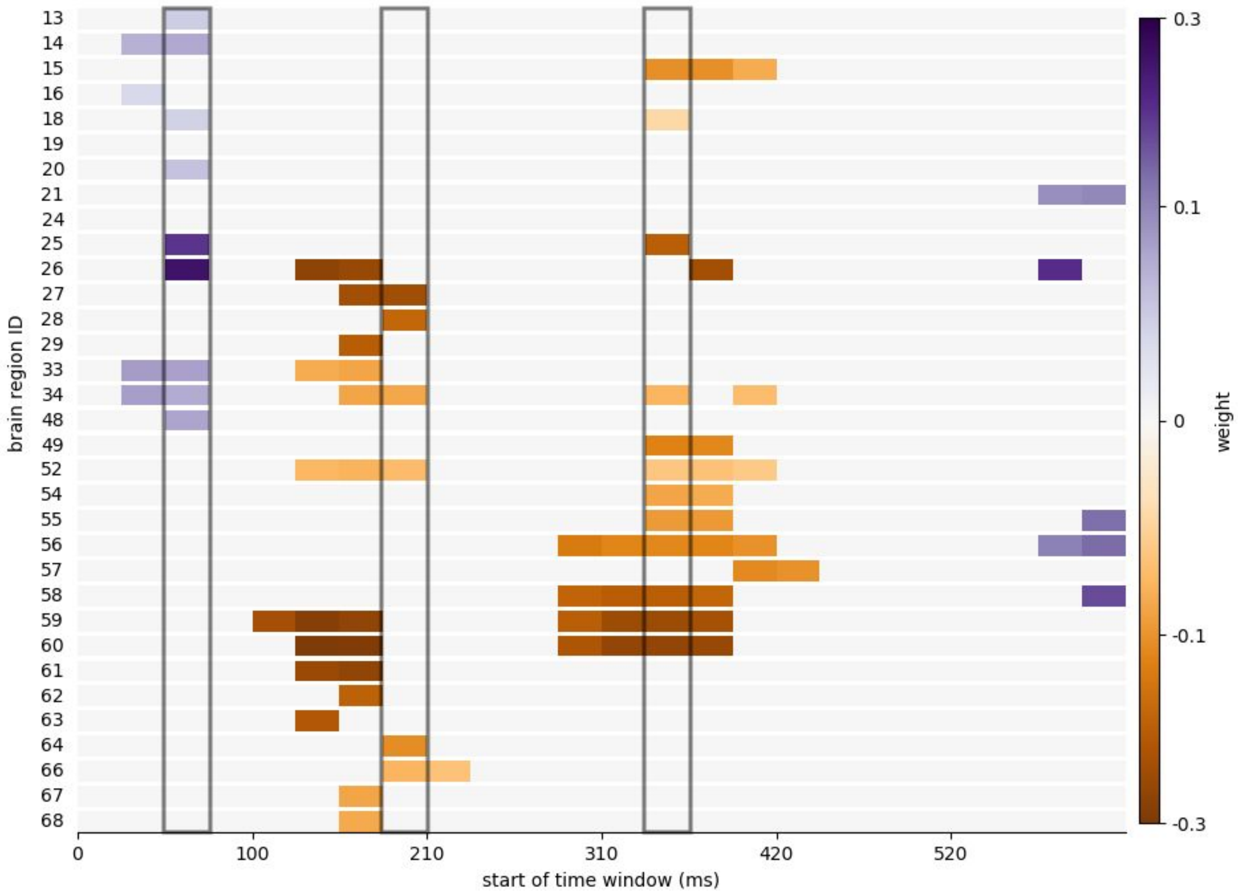


Figure S4: (related to Figure 4) Time courses of brain regions that are significantly different between faces and scrambled conditions. Gray boxes mark time windows for which surface renderings are shown in Figure 4. See Figure S3 for a legend of the brain regions' locations.
